# Estimating cis and trans contributions to differences in gene regulation

**DOI:** 10.1101/2024.07.13.603403

**Authors:** Ingileif B. Hallgrímsdóttir, Maria Carilli, Lior Pachter

**Affiliations:** Division of Biology and Biological Engineering, California Institute of Technology, Pasadena, CA, 91125, USA; Department of Computing and Mathematical Sciences, California Institute of Technology, Pasadena, CA, 91125, USA

## Abstract

We describe a coordinate system and associated hypothesis testing framework for determining whether *cis* or *trans* regulation is responsible for differences in gene expression between two homozygous strains or species. We apply our framework to data from single replicate studies on yeast strains and human-chimpanzee hybrid cells, as well as to data from a mouse study with replicates, showing marked differences between our gene regulatory assignments and those previously reported. We also show how our multi-sample framework can determine the context dependency of *cis* and *trans* effects as well as explicitly model different hypotheses regarding the underlying mechanism of *trans* regulation.

## Introduction

In 1961, Jacob and Monod developed a theory of gene regulation in which they distinguished local effects (*cis*) from distal regulation (*trans*) (1). Their work immediately raised the question of the relative contributions of these two regulation modalities (2, 3). One approach for assessing whether *cis* or *trans* regulation is responsible for differences in gene expression between strains or species is to compare differences in expression of genes in parents to allele-specific differences in *F*_1_ hybrids. This approach was explored in (4), who used crosses of C57BL/6J and CAST/Ei mice to study regulatory mechanisms that could explain differences in gene expression between parental strains. Their approach was developed in (5, 6), who used pyrosequencing to study the regulation differences between *D. melanogaster* and *D. simulans*. With the advent of RNA-seq, genome-wide scans were possible, and (7) examined RNA-seq from *F*_1_ crosses of C57BL/6J and CAST/EiJ to tease apart *cis* and *trans* contributions to gene expression differences between the parental strains. Similarly, (8) performed such an RNA-seq analysis using *Drosophila* lines.

Formally, the idea of using crosses to study *cis* and *trans* contributions to differences in gene expression between strains or species is as follows: consider a gene with expression *X*_*P* 1_ in a homozygous strain 1, *X*_*P* 2_ in a homozygous strain 2, and expression *X*_*H*1_ for the haplotype from strain 1 in the *F*_1_ cross of 1 and 2, and expression *X*_*H*2_ for the haplotype from strain 2 in the *F*_1_ cross of 1 and 2. Let 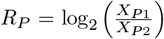 and 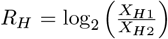. That is, *R_P_* corresponds to the log2 fold change difference in expression between the two parental strains, and *R*_*H*_ to the log2 fold change difference between the expression of the hybrid hap-lotypes in the *F*_1_ offspring. The connection between *R*_*P*_, *R*_*H*_, and regulation is as follows: consider that a gene can be regulated via *cis, trans*, or both (Fig. 1A). Gene expression measurements in the parents and hybrid (Fig. 1B), can be used to infer the nature of regulation underlying the difference in expression in the parental strains (Fig. 1C). Specifically, amending the classification of (6), we have:

- ***conserved:*** No change in gene expression indicating there has been no change in regulation, i.e. *R*_*P*_ = 0 and *R*_*H*_ = 0, which implies *R*_*P*_ − *R*_*H*_ = 0.
- ***cis:*** The relative difference in gene expression between the parents is the same as between the haplotypes in the hybrid indicating that the difference in parents is due to local *cis* effects, i.e. *R*_*H*_ ≠ 0, *R*_*P*_ 0, and *R*_*P*_ = *R*_*H*_ which implies *R*_*P*_ −*R*_*H*_ = 0, arises from changes only in cis-regulatory elements.
- ***trans:*** Gene expression from the two haplotypes in the hybrid is the same, indicating that differences between the parents resulted from non-local *trans* regulation, i.e. *R*_*H*_ = 0 and *R*_*P*_ 0, which implies *R*_*P*_ − *R*_*H*_ ≠ 0, arises from a change only in *trans*-regulatory elements.
- ***cis + trans:*** *R*_*H*_ 0 and *R*_*P*_ *≠ R*_*H*_ with sgn(*R*_*H*_) = sgn(*R*_*P*_ −*R*_*H*_) arises as a result of change in both *cis*- and *trans*-regulatory elements with changes in *cis* and *trans* contributing to changes in gene expression between strains in the same direction.
- ***cis*** × ***trans:*** *R*_*H*_ 0 and *R*_*P*_ *R*_*H*_ with sgn(*R*_*H*_) ≠ sgn(*R*_*P*_ −*R*_*H*_) arises as a result of compensatory change in both *cis*- and *trans*-regulatory elements with changes in *cis* and *trans* contributing to changes in gene expression between strains in the opposite direction.

**Fig. 1.**
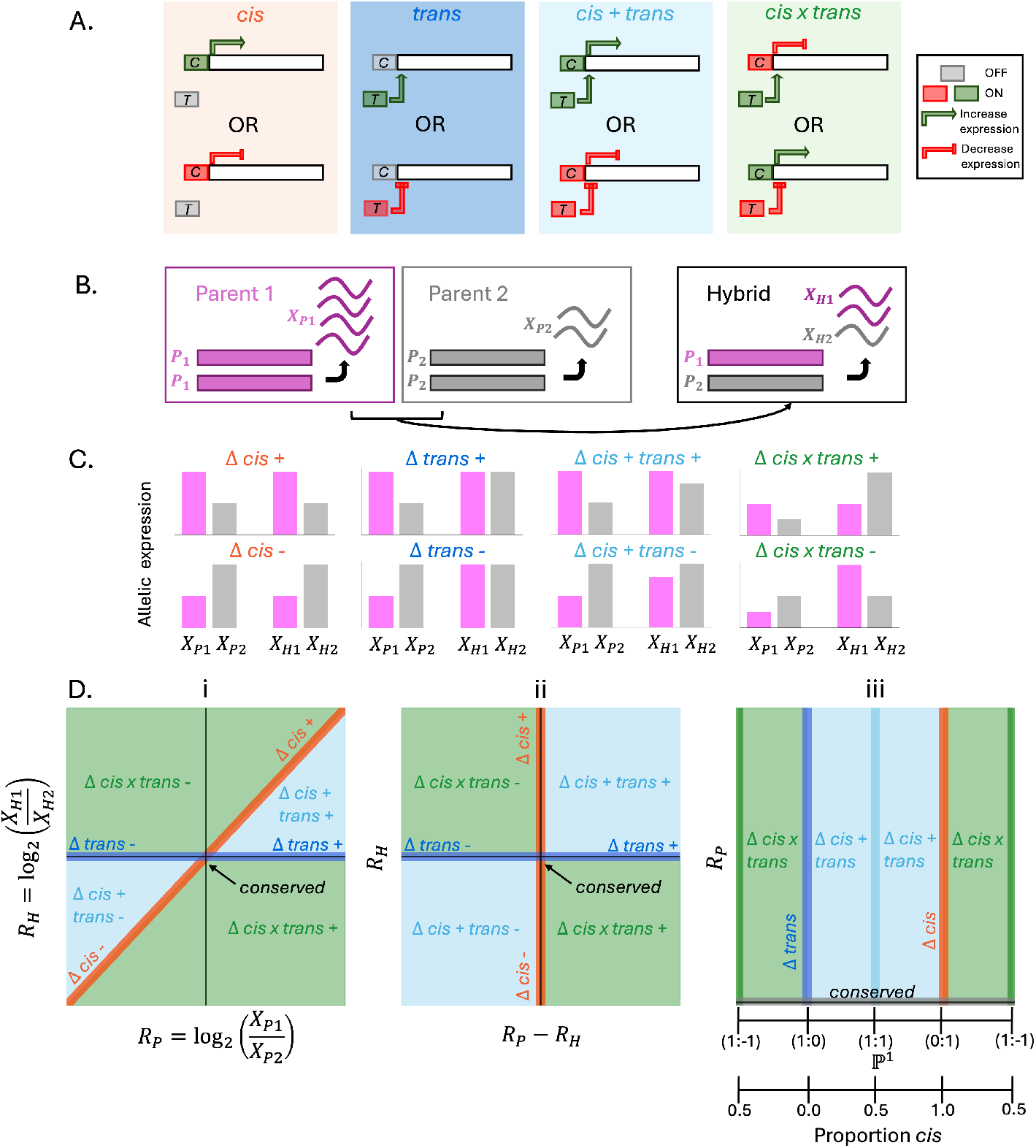
Geometry of parental and hybrid expression ratios can be used to assess regulatory differences between homozygous crosses. A) Diagram of the *cis, trans, cis + trans*, and *cis* × *trans* types of regulatory differences. B) Homozygous parents with haplotypes *P*_1_ and *P*_2_, counts of RNA molecules for a gene (*X*_*P* 1_ and *X*_*P* 2_ respectively), and the haplotypes in an *F*_1_ hybrid along with counts for the gene (*X*_*H*1_ and *X*_*H*2_). C) Differences in regulation between the parents are reflected in distinct ratios between counts *X*_*P*1_, *X*_*P* 2_ and *X*_*H*1_, *X*_*H*2_. D. i) Illustration of how regulation differences emerge in log-fold changes *R*_*P*_ and *R*_*H*_, D. ii) Linear transformation of 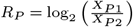 and 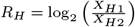 to yield orthogonal *cis* and *trans* coordinates, D. iii) Representation of proportion *cis* in real projective space ℙ^1^.

This classification corrects (6), which fails to properly assign regulation differences to *cis + trans* when both *R*_*H*_ < 0 and *R*_*P*_ < 0. The relationships between *R*_*P*_ and *R*_*H*_ in general, can be visualized as lines and regions in a 2D plot (5), as illustrated in Fig. 1D(i). While this direct representation of *R*_*P*_ and *R*_*H*_ is useful, a quantitative assessment of the gene regulatory modalities reflected in *R*_*P*_ and *R*_*H*_ requires a biologically meaningful notion of distance between points in Fig. 1D(i). Consider, for example, the situation where *X*_*P* 1_ = 2*X*_*P* 2_ and *X*_*H*1_ = 2*X*_*H*2_, i.e. a 2-fold change in gene expression between the parents due solely to *cis* regulation which corresponds to the point (1, 1) in Fig. 1D(i). The distance from this point to the origin (conserved), is 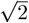, whereas the same 2-fold difference in gene expression in the parents due solely to *trans* regulation with *R*_*P*_ = 1 and *R*_*H*_ = 0 is distance 1 from the origin. This imbalance can be corrected via a linear transformation.

## Results

### Geometry

To decouple the effects of *cis* and *trans* regulation on *R*_*P*_ and *R*_*H*_ we begin by noting that if the difference in parental expression is solely due to *cis* regulation, then *R*_*P*_ = *R*_*H*_, or equivalently *R*_*P*_ − *R*_*H*_ = 0 (orange vertical line in Fig. 1D(ii)). If the difference in parental expression is solely due to *trans* regulation, then *R*_*H*_ = 0 (blue horizontal line in Fig. 1D(ii)). Therefore, the transformation from Fig. 1D(ii) to Fig. 1D(i) is obtained by

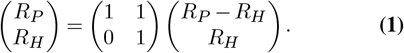

Thus, the inverse transformation from the coordinate system in Fig. 1D(i) to the coordinate system in Fig. 1D(ii) is given by

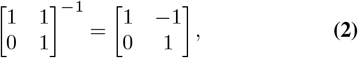

i.e., the transformation inverts the mapping of the vertical axis to the diagonal *cis* line.

The determination of whether a difference in parental gene expression is due to *cis* or *trans* can now be understood to be an assessment of whether the line passing through the origin and a point (*R*_*P*_ −*R*_*H*_, *R*_*H*_) is a perturbation (due to noise in gene expression measurement) of the line Δ *trans*, the line Δ *cis* or sufficiently far away from the axes in Fig. 1D(ii) to merit a designation of Δ *cis + trans* or Δ *cis* × *trans* (the designations are enumerated in Supp. Table 1). In other words, the sufficient statistic is a point in real projective space ℙ^1^ (Fig. 1D(iii)), and the proportion of the difference in gene expression between parents that can be attributed to *cis* can be understood to be a scaling of the angle of the line through the origin corresponding to the point in ℙ^1^, i.e.,

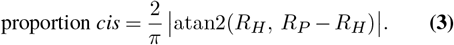

### Statistics

The determination of whether a measurement (*R*_*P*_, *R*_*H*_) reflects a difference in gene expression between parents due to *cis* or *trans* regulation, or both, requires a statistical assessment (10). Specifically, hypothesis tests can be used to reject a null hypothesis of a difference in gene ex-pression being due solely to *trans* or solely to *cis*. Geometrically, as evident from the linear transformation on log-fold changes, these two tests correspond to testing whether one can reject the null hypotheses that (*R*_*P*_ −*R*_*H*_, *R*_*H*_) is located on the *x*- and *y*-axes respectively.

We first developed a framework for hypothesis testing in this case where there are no replicates of the gene expression measurements, as in (9), which consists of bulk RNA-seq performed on hybrid and parental strains of genetically divergent *Saccharomyces cerevisiae* (see Methods, Supp. Figs. 1,2). Briefly, (9) generated hybrid crosses from 26 parental yeast isolates derived from diverse environmental conditions. In (9), genes were classified according to the representation shown in Fig. 1D(i). First, both *R*_*P*_ and *R*_*H*_ were tested for statistically significant differences from 0 and assigned “null” (our *conserved*) if both tests failed to reject the null hypothesis (one-sample allele-specific expression tests); then, changes between parental and hybrid allelic ratios were tested for significance (two-sample allele-specific expression tests). For genes that passed significance thresholds, regulatory assignments were made by dividing the untransformed 2D plane into cones (Fig. 2A). We reassigned genes based on our hypothesis tests as derived from the transformed coordi-nate system (Fig. 2C), with one test with the null hypothesis of *cis* regulation and one with the null hypothesis of *trans* regulation, thereby putting the two regulation strategies on equal footing. Compared to the original study, we found major differences in assignment (Fig. 2B, Supp. Fig. 3).

**Fig. 2.**
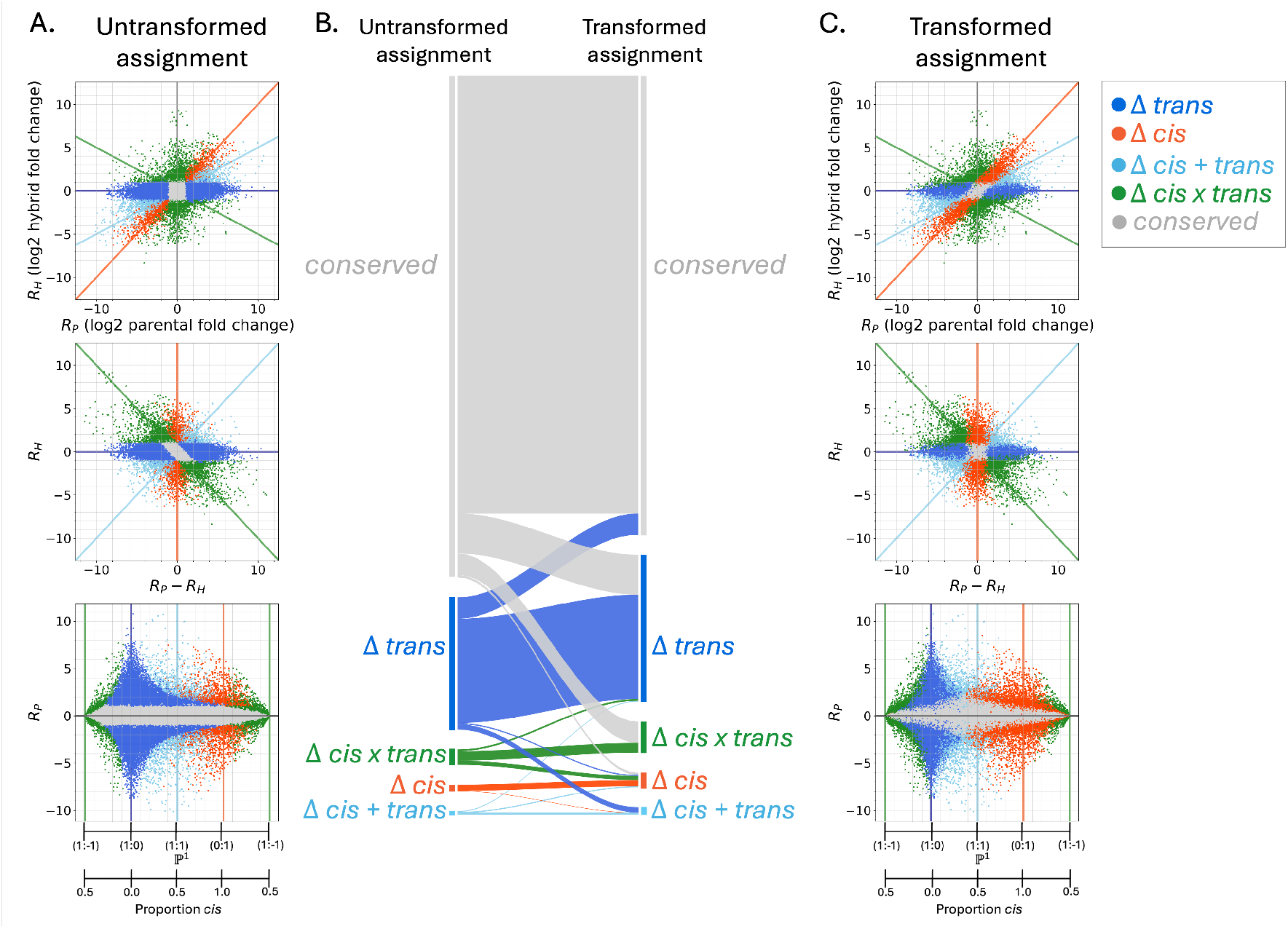
Contributions of *cis* and *trans* regulation to differences in gene expression between yeast strains, with data from (9) (n = 285,777 gene-cross combinations from 179 unique yeast parent crosses). A) Untransformed ratios, transformed ratios, and log2 of parental fold change versus proportion *cis*, colored by regulatory assignments from (9). B) Comparison of the results of (9) to the assignment of genes based on the proposed hypothesis testing and geometric assignment procedure. C) Reanalysis of the 285,777 gene-cross expression data from (9) using the proposed hypothesis testing and geometric assignment procedure.

**Fig. 3.**
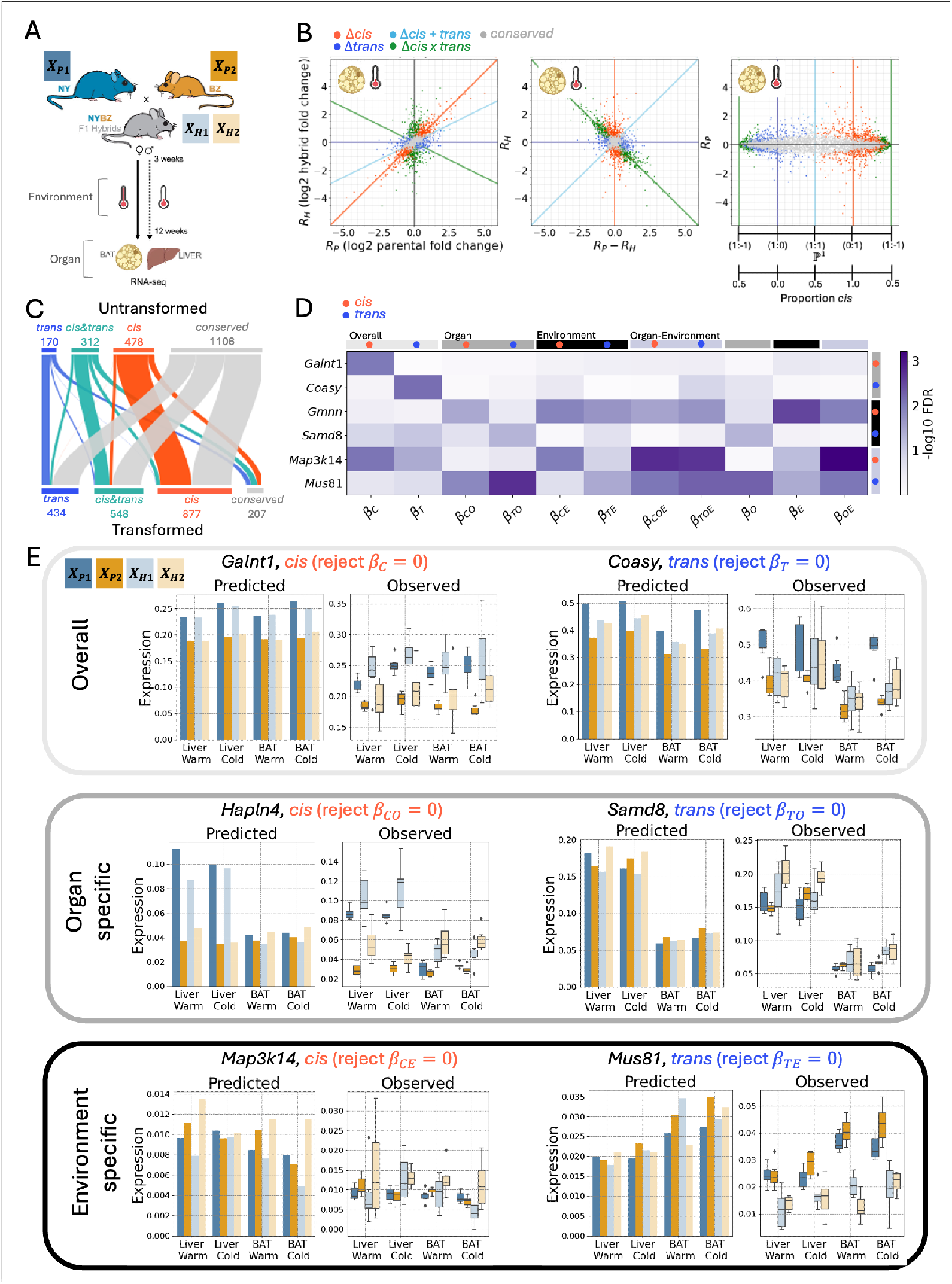
Multi-sample regulatory assignment can be determined using generalized linear models and condition-specific hypothesis tests. A) Cartoon of experimental design set up from original study (adapted from Fig. 1C of (13)). B) Model predicted ratios colored by GLM determined regulatory assignments as described in Methods. The log additive model was fit to brown adipose tissue from male mice reared in the cold (n = 6 NY mice, n = 6 BZ mice, and n = 6 hybrid crosses, and n = 5,970 genes). C) Untransformed (original) regulatory assignments (reported in (13)) derived from three independent model fits and statistical tests differ from transformed regulatory assignments derived using the unified GLM framework (genes and assignments from B, excepting genes that were categorized as conserved by both the original and our testing procedure, which are not shown). D) The log additive model was fit to samples from all four environment/organ pairs (BAT/cold, BAT/warm, liver/cold, liver/warm, n = 6 NY/BZ/hybrid samples per condition) with weights for environment and organ-specific regulatory effects. The negative log10 of false discovery rates (FDR) highlights several genes that exhibit significant *cis* or *trans* behavior overall or only in an organ or environment-specific manner. E) For the same genes, GLM predicted (bar plots) and observed gene expression (box plots with counts normalized by the total count per sample for parents and per haplotype for hybrids) for parental and hybrid alleles (n = 6 samples per box, with the box showing values’ interquartile ranges and a line at the mean and whiskers extending to min/max values).

Interestingly, whereas (9) conclude that “the transcriptome is globally buffered at the genetic level mainly due to trans-regulatory variation in the population”, we find that a considerable amount of difference in gene expression can be attributed to *cis* (Supp. Fig. 3C), with the difference due to (9) deriving assignments in the untransformed coordinate system using a statistical testing procedure that treats *cis* and *trans* regulation asymmetrically. Specifically (9) report 57,253 cases where gene expression difference is due to *trans* regulation (Supp. Fig. 3A). We find 51,627 cases that can be assigned to *trans* (Supp. Fig. 3B). These numbers are similar; however, (9) report 2,804 cases assigned to *cis* (Supp. Fig. 3A), whereas we find *17,112* (Supp. Fig. 3B). Furthermore, we find 4,807 cases assigned to *cis + trans* (Supp. Fig. 3B) versus 1,727 in (9) (Supp. Fig. 3A). Overall, there is a marked difference between our results and those of (9) (Supp. Fig. 3C).

In the case where replicates have been obtained, a generalized linear model (GLM), as is commonly used for differential expression in bulk RNA-seq (11, 12), can be adapted to utilize gene expression variance estimates within parents and hybrids. We tested our approach (see Methods) on data from (13), in which two wild-derived mouse strains and their F1 hybrids were used to study the effect of genotype and environment (temperature) on gene expression divergence in rapidly evolving strains. Two inbred lines of house mice were derived from mice from a cold environment (Saratoga Springs, New York, USA) and a warm environment (Manaus, Amazonas, Brazil) (13), and two tissues, brown adipose tissue (BAT) and liver, were evaluated in each of the strains and their F1 hybrids, also raised at the two different temperatures. The experiment was performed in six replicates each of male and female parents, as well as six replicates each of male and female F1 hybrids (13) (Fig. 3A). The GLM framework allows the joint analysis of all samples (parents and hybrids) rather than requiring that separate tests be run on separate subsets of the data (e.g., differential expression between parents, differential expression between hybrid allelic expression, and a ratio test as in (13)). It further enables explicit modeling of specific biological hypotheses of *trans* regulatory action (see Methods, Supp. Figs. 4 – 6).

**Fig. 4.**
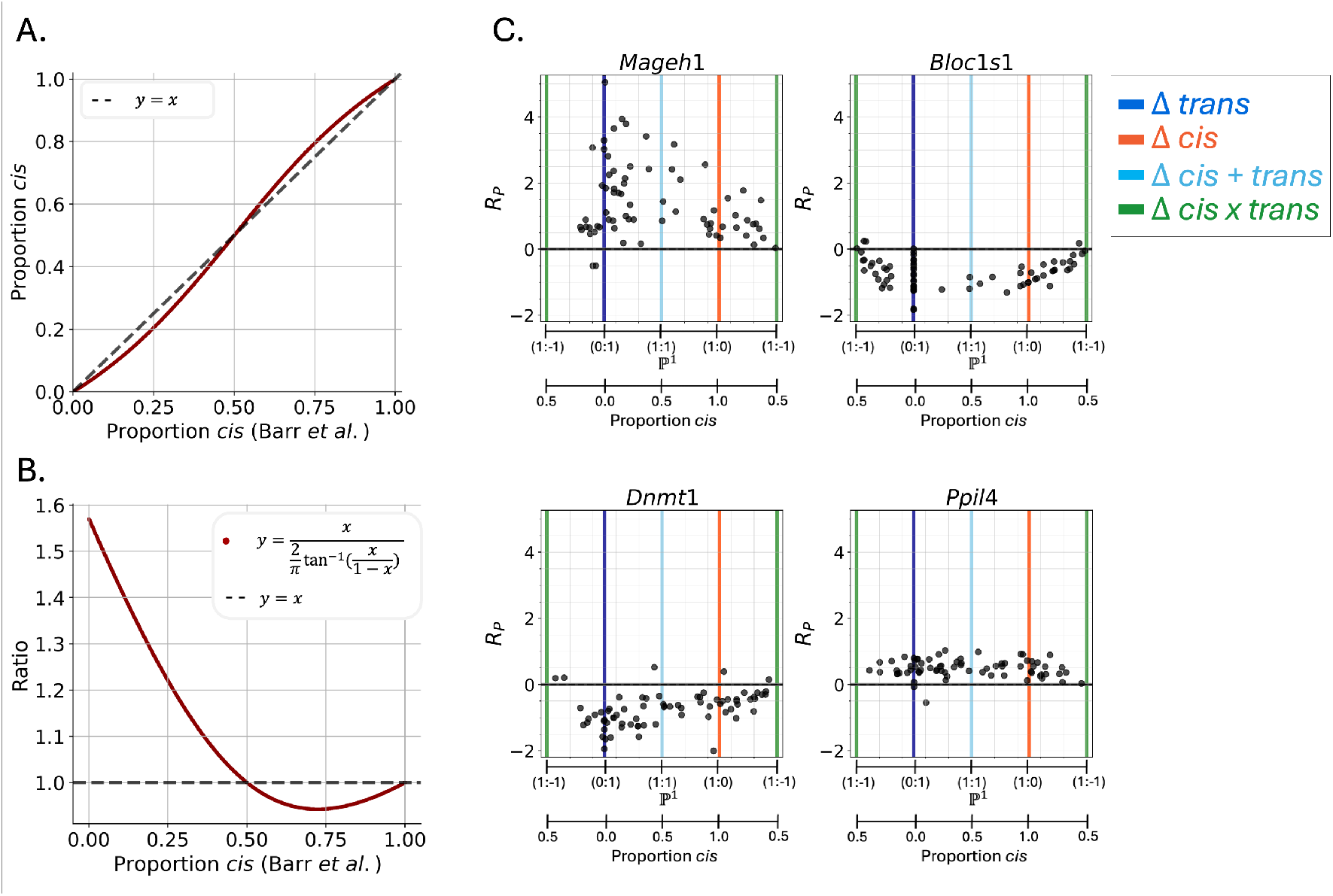
In addition to broad regulatory categories, genes can display subtle differences in their proportion of *cis* regulation. A) Comparison of slope (as previously defined (17), x-axis) and angle (y-axis) for determining proportion *cis*. B) Ratio of slope and angle determination of proportion *cis* (dark red line, dashed line displays equality). Note that at small values of proportion *cis*, the slope can be over 50% the determined angle. C) Gene expression data from (17) (human-chimpanzee crossed cell lines) was used to determine cell type specific proportion *cis*. These four genes display high variance in proportion *cis* across cell types (black points are determined proportion *cis* in n = 72 cell types).

Restricting our analysis to a single tissue, temperature, and sex (brown adipose tissue collected from male mice reared in the cold temperature), we fit a GLM with the log additive hypothesis and classified genes based on the reduced models of no *cis* or no *trans* regulation (see Methods, Fig. 3B). Consistent with our single sample analyses, we noted an increase in the number of *cis* assigned genes from 478 in the original study (13) to 877 (Fig. 3C). This trend held in different tissue and temperatures (Supp. Fig. 7). We further fit the log additive model to all male samples, including environment (cold/warm) and organ (liver/BAT) specific *cis* and *trans* weights (Fig. 3D). We identified genes that exhibited significant overall *cis* and *trans* regulatory changes (Fig. 3E, top panel), as well as those that were only significantly different in one organ (Fig. 3E, middle panel) or environment (Fig. 3E, bottom panel). Among these genes were several with functions related to body fat and temperature. For example, *Coasy*, which was identified as having an overall significant *trans* regulatory difference between the strains (Fig. 3E, top panel), codes for CoA (coenzyme A) synthase. CoA is critical for energy production and fatty acid synthesis, and other proteins involved in its production pathway have been shown to regulate adaptive thermogenesis in mice (14). *Samd8*, identified as having tissue-specific regulatory patterns, codes for a protein that synthesizes sphingolipids, which act as key regulators in brown adipose tissue with accumulation linked to tissue aging and impaired lipid homeostasis (15). The gene that produces mitogen-activated protein kinase NIK, *Map3k14*, was found to have a significant temperature-dependent *cis* effect: related proteins in the MAPK family mediate heat-shock reactions in mice, with their inhibition leading to reduced protection against heat (16). These genes are plausible candidates for differential regulation between mouse strains adapting to life in cold and warm environments. The flexibility of the GLM framework facilitates identification of genes with unique regulatory patterns, such as the tissue and organ-specific effects, a key improvement over post-hoc comparisons of multiple model fits that would be otherwise required.

### Proportion *cis*

In addition to naturally revealing the appropriate hypothesis tests to conduct for attribution of gene expression difference in parents to *cis* or *trans*, the linear transformation we propose leads directly to a meaningful measure of the proportion of difference in gene expression that can be attributed to *cis* regulation (or *trans*) (Equation 3). To illustrate this, we re-analyzed a dataset of gene expression from human-chimpanzee cell line hybrids (17), calculating the proportion *cis* according to Equation 3. In (17), the proportion *cis* was calculated using slope, as is natural to do when working in the untransformed coordinate framework, whereas the correct calculation (Equation 4) uses angle in the transformed coordinate system. While the absolute differences are small (Fig. 4A), with a maximum difference of 0.045 (see Methods), the relative difference is large when the proportion *cis* is small (Fig. 4B), and can be as high as 57% (see Methods). Moreover, our computation provides a biologically interpretable measure of proportion *cis*. We found several genes in (17) exhibiting high variance in proportion *cis* (Fig. 4C), and identified interesting differences between cell types (Supp. Fig. 8 – 11).

## Discussion

The use of crosses between strains to identify the nature of differential regulation is a powerful tool for genetics studies that is particularly relevant now that single-cell RNA-seq can be used for cell type resolution. Moreover, while original studies were limited to a handful of genes, genome-wide single-cell RNA-seq assays can complement genome-wide eQTL studies.

We have shown that geometric considerations reveal the need for applying a linear transformation prior to visualization. The linear transformation highlights independent axes that lead naturally to hypothesis tests for classifying genes according to the type of regulation underlying differences in gene expression between parents. While we have focused on explaining differences between parents that fall into five categories (conserved, *cis, trans, cis + trans, cis*× *trans*), our approach can be extended to finer classifications such as in (9). We note that in the hypothesis testing framework, referring to a gene as having differences in expression explained by *cis* or *trans* is technically incorrect. This is because the rejection of the *cis* null hypothesis only shows that the difference in gene expression is not due solely to *cis* regulation. This does not mean that the difference in gene expression can or should be attributed solely to *trans*. The same is the case for rejecting the *trans* hypothesis. In Fig. 2, our coloring of genes as *cis* or *trans* is therefore not precise, but we have done so to facilitate comparisons to previous work.

Our single sample statistical tests depend on the assumption that read counts are binomially distributed, which is standard in the absence of biological replicates. However, when replicate samples are available, we can better assess the extent of technical variation and use generalized linear models. Although our results are shown using commonly applied estimation techniques for negative binomially distributed counts (11), we also measure the extent to which differing estimates of variation change the results (Supplement) (11, 12). We encourage experimentalists to include enough samples in the experimental design such that heuristic estimations are unnecessary and the observed sample variation can be used for hypothesis testing.

Finally, we note that our framework is general and can be applied to more complex experimental designs, regulatory hypotheses, and phenotypes other than gene expression. For example, the versatility of our GLM framework allows modeling of *cis* or *trans* differences in specific conditions and in interactions between specific conditions. In terms of additional phenotypes, with single-cell RNA sequence data our approach could be used in conjunction with methods such as (18, 19) to assess the regulation mechanisms underlying differences in biophysical aspects of gene transcription, splicing and degradation. Such extensions will be particularly interesting to explore in conjunction with complementary modalities (20, 21).

## Data and Code Availability

All code to download data and generate the main and supplementary figures is available at https://github.com/pachterlab/HCP_2024, and an R package (*XgeneR*) to implement the multi-sample tests at.

## Acknowledgements

IH and LP were funded, in part, by NIH 5UM1HG012077-02.MC was funded by a NSF graduate research fellowship under Grant No. 2139433. This work was additionally supported by the Caltech Bioinformatics Resource Center.

## Methods

### Acquisition and preprocessing of data from yeast crosses

Processed RNA-seq data from (9) were obtained from http://1002genomes.u-strasbg.fr/files/Diallel_RNAseq/ASE in the file ‘Datafile2_ase_sum_20230609.tab.’ This file included reported values for allelic expression in parents and expression per parental allele in hybrids for 179 unique parent-hybrid trios and 285,777 unique gene-trio combinations. Details on the estimation of allelic expression are described in the original study (9). As integer value counts are required for the binomial and the two sample binomial ratio statistical tests, reported values were rounded to the nearest integer before statistical tests were performed.

In (9), 7 categories were considered (*reverse, attenuating, reinforcing, compensatory, cis only, trans only* and *null*). In order to maintain consistency with (6), we collapsed these into four categories: *cis, trans, cis + trans*, and *cis* × *trans* as follows: reverse → *cis* × *trans, cis only* → *cis, trans only* → *trans*, null → conserved, and attenuating, reinforcing and compensatory → *cis* × *trans* if sgn(*R*_*P*_ − *R*_*H*_) = sgn(*R*_*H*_) and *cis* + *trans* otherwise.

### Acquisition of data from human-chimpanzee crosses

The reported log2 fold changes between human and chimpanzee parental gene expression (*R*_*P*_) and hybrid allelic log2 fold changes for human-chimpanzee hybrid cell lines (*R*_*H*_) were obtained from (17) for 72 cell types and 14,487 genes per cell type. The reported log2 fold changes were used without modification to recalculate proportion *cis* as

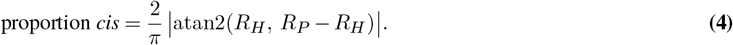

The provided data also included reported proportion *cis* per gene per cell type, defined as

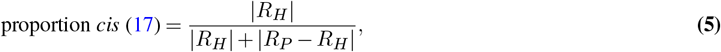

which were directly compared to Equation 4.

The maximum difference between 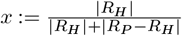and 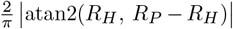 is 0.045 at *x* = 0.239 and *x* = 0.761. The maximum of the ratio is 1.57 and occurs at *x ≈* 0.00006 (Fig. 3B).

### Acquisition of data from mouse crosses

(13) performed a study to assess regulatory strategies driving environmental adaptations in house mouse from two different climates. To do so, they created two inbred lines derived from wild house mice from Saratoga Springs, New York, USA (NY, cold climate) and Manaus, Amazonas, Brazil (BZ, warm climate) and crossed them. Mice from both inbred lines and their hybrid crosses were housed in warm (21 °C) and cold (5 °C) conditions, then liver and brown adipose tissue (BAT) samples from male and female mice were collected. RNA sequencing, alignment, and allele-specific expression (ASE) quantification for parental and hybrid tissue samples were performed as described in the original publication ((13)).

To assign genes’ regulatory categories, they performed three statistical tests on the parental counts and ASE in the hybrids (using *DESeq2* (12)): first, they tested for differential expression between lines using only parental counts; second, they tested for differential expression between alleles in the hybrids using only hybrid ASE; and third, they tested for ratio differences between alleles in the parents and alleles in the hybrids using parental counts and ASE (details in the Methods of (13)). They assigned genes as *cis, trans, cis & trans* or *conserved or ambiguous* based on the outcomes of the three tests, as described in (13). For consistent comparison, we collapse the categories *cis + trans* and *cis x trans* to *cis & trans*.

We obtained parental counts, regulatory assignments for genes in the four separate environment (warm/cold) and organ (liver/BAT) combinations, and sample metadata from the GitHub repository https://github.com/malballinger/BallingerMack_PNAS_2023/. We obtained ASE for hybrids from the authors directly. We applied our regulatory assignment framework (see the subsection ***Generalized linear models for multiple samples***) to male mice (for each of the four environment-organ combination there were 6 SARA mice, 6 MANA mice, and 6 hybrids, for a total of 18 mice and 24 rows in the design matrix, as each hybrid contributes expression levels for two alleles) for each environment/organ condition separately (liver/warm, liver/cold, BAT/warm, BAT/cold); and for all male samples together (total of 72 mice and 96 rows of the design matrix). We used the log-additive *trans* hypothesis model for testing all samples and condition-specific effects and for (liver/warm, liver/cold, and BAT/warm) male samples. For all warm samples, there were 5,897 fit genes, and 5,970 for cold samples. We fit the log additive, dominant, and free models to the BAT/cold male samples, with comparisons of fit weights, significance values, and gene regulatory assignments shown in Supp. Figs. 4 – 6. All fits were implemented using the XgeneR package with edgeR version 4.0.16 (11).

## Statistical tests

### Binomial test

To test the null hypothesis that there is no difference between expression of parental alleles in hybrids (or that the difference in regulation is purely *trans*) was performed on rounded integer counts from (9) using the function `scipy.stats.binomtest` (22). This tests for the probability of having observed a value at least as extreme as *k* successes given probability *p* of success and a total of *N* trials by summing over binomial probabilities:

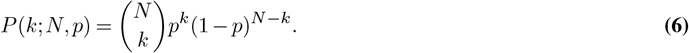

If *X*_*H*1_ is the allelic expression in the hybrid of one parental allele and *X*_*H*2_ is the allelic expression in the hybrid of the other parental allele, the binomial test was performed with *k* = *X*_*H*1_, *N* = *X*_*H*1_ + *X*_*H*2_, *p* = 0.5 and a two-sided alternative hypothesis.

### Two sample binomial ratio test

To test the null hypothesis that there is no difference in the ratio of expression of alleles in parents and allelic expression in hybrids (or that the difference in regulation between parents is purely *cis*), we performed a two sample binomial ratio test in which we assumed both *X*_*P* 1_ and *X*_*H*1_ are sampled from a binomial distribution with the same probability of success *p*_*s*_. Using

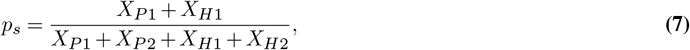

we calculated the grid of probabilities over possible *P*_1_ and *H*_1_ values:

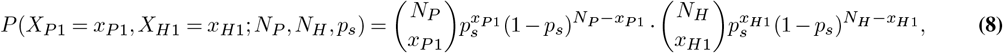

where *N*_*P*_ = *X*_*P* 1_ + *X*_*P* 2_ is the total number of counts from parents and *N*_*H*_ = *X*_*H*1_ + *X*_*H*2_ is the total number of counts from the hybrid. We then calculated the probability of having observed values at least as extreme as the observed *X*_*P* 1_ and *X*_*H*1_.

To account for multiple testing, we corrected both binomial and two sample binomial significance values using the Benjamini-Hochberg correction for false discovery rates (23). We rejected the null hypothesis for tests with false discovery rates less than 0.05.

### Generalized linear models for multiple samples

We propose a generalized linear model (GLM) framework for testing the null hypotheses of *no cis* and *no trans* regulation, both overall and condition-specific, when multiple samples per condition are available. Our framework differs from previous applications of GLMs for *cis/trans* distinction (13) by virtue of using one GLM for all data (parental counts, hybrid ASE values, with all conditions represented), rather than fitting three separate models to different subsets of the data (e.g., only parental counts for one test or only hybrid ASE values for another). We do this by developing a model with weights for overall and condition-specific *cis* and *trans* regulatory effects, as well as including condition intercepts to properly attribute observed allele expression patterns to regulation or condition. In addition, our approach allows us to propose different hypotheses about the nature of the *trans* regulatory effect: if it is log-additive, dominant, or free (described below). To properly account for the overdispersion and discreteness of RNA-seq counts, we fit a GLM with a negative binomial likelihood function and log link function.

#### One condition

In the simplest case, we set expression of the first parental allele undergoing only *cis* regulation to be the intercept *β* and include weights for the *cis* regulatory difference (*β*_*C*_) and *trans* regulatory differences (*β*_*T*_) between parental strains. This allows the normalized expression from the parental alleles (*X*_*P* 1_, *X*_*P* 2_) and normalized allelic expression from the hybrids (*X*_*H*1_, *X*_*H*2_) to be modeled under three different hypotheses of *trans* regulation.

The first hypothesis (**log-additive**) assumes that the *trans* expression difference between parental strains at each allele is influenced by both sets of parental chromosomes multiplicatively, and is thus reduced to the square root of the full parental effect in the hybrids. The design matrix is accordingly

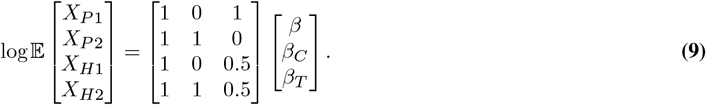

A second hypothesis (**dominant**) assumes that the *trans* expression difference between parental strains is the same regardless of if both sets or one set of parental chromosomes are present: it is the same in the parents and hybrids, and the design matrix is

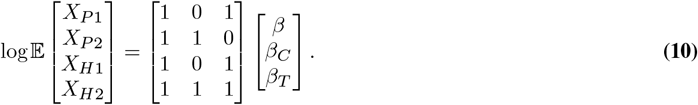

Finally, and most analogous to previous hypothesis testing methods (13), *trans* expression differences between parental strains can have an unconstrained relationship (**free**) with their effects in the hybrids by allowing each hybrid to have individual specific effects (denoted below with the subscript *i* indexing the hybrid samples). The resulting design matrix is

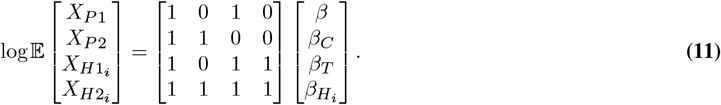

For all models, we can test the null hypothesis of *no cis* differences between parental strains by testing if *β*_*C*_ = 0 and the null hypothesis of *no trans* differences by testing if *β*_*T*_ = 0.

#### Multiple conditions

As more conditions are added to the experimental design, they can be naturally incorporated into the GLM framework, with condition-specific *cis* and *trans* weights as well as intercepts and interaction terms. For example, we fit data for liver and BAT samples for mice grown in warm and cold environments (four separate organ/environment) using the following model under the log-additive *trans* hypothesis:

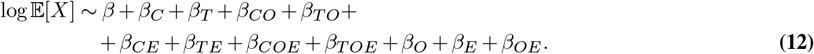

The subscripts *C, T, O* and *E* refer respectively to differential *cis* regulation between strains, differential *trans* regulation between strains, organ, and environment effects, with combinations denoting their interactions (e.g., *β*_*CO*_ models the organ-specific *cis* effect). This is also compatible with any of the three previously described *trans* hypotheses (log-additive, dominant, and free).

#### Implementations

We implemented GLMs for the one condition in edgeR version 4.0.16 (11) and DESeq2 version 1.42.1 (12), showing differences in estimated gene variance, data likelihoods and hypothesis test significance values in Supp. Fig. 12. In particular, we tested significance using a likelihood ratio test (LRT) between the full and reduced models. Differences in significance values lead to differences in regulatory assignments, especially pertinent in this our case in which both *no cis* and *no trans* hypothesis testing results are important for regulatory categorization. As edgeR is the more conservative method, estimating higher variance and false discovery rates (Supp. Fig. 12), we used it for full model fits and regulatory assignments. We corrected LRT p-values for each hypothesis test for the number of genes tested using the Benjamini-Hochberg correction (23) to obtain false discovery rates, and rejected null hypotheses with false discovery rates less than 0.05.

### Classifying genes based on statistical test results

The results of the two tests for single sample experiments or experiments with replicates are used to categorize genes as outlined in Table 1.

**Table 1.** Statistical assignments of gene regulatory differences. Genes assigned to *cis, trans, cis + trans* or *cis* × *trans* regulation based on two statistical tests, described above for single sample or replicate experiments. If both statistical tests are rejected, regulatory assignment can be made by locating the point (*R*_*P*_ − *R*_*H*_, *R*_*H*_) (see Results Section, Geometry) in the transformed space and following the geometric assignments listed in Supp. Table 1.

| Single sample: Binomial test<br>Replicates: $\beta_C = 0$<br>(null: *no cis*) | Single sample: Binomial ratio test<br>Replicates: $\beta_T = 0$<br>(null: *no trans*) | Classification |
| --- | --- | --- |
| Fail to reject | Fail to reject | conserved |
| Reject | Fail to reject | *cis* |
| Fail to reject | Reject | *trans* |
| Reject | Reject | *cis & trans* (*cis* + *trans* or *cis* × *trans* by geometry) |

Note that some care must be taken when interpreting a *cis* or *trans* assignment within this hypothesis testing framework. For example, a *cis* assignment is made when “no *cis*” can be rejected but “no *trans*” cannot be rejected. Failure to reject “no *trans*” does not imply that one can accept the null hypothesis, but we utilize the assignment of *cis* in this case for simplicity for users. However, this caveat should be taken into account when interpreting the assignments of *cis, trans*, and *cis & trans* assignments.

