## Supplement for "Estimating cis and trans contributions to differences in gene regulation"

#### Supplementary tables

| $R_P - R_H$ | $R_P$ | $R_H$ | **Regulatory assignment** |
|---|---|---|---|
| 0 | 0 | 0 | conserved |
| 0 | + | + | Δ *cis* + |
| 0 | - | - | Δ *cis* - |
| + | + | 0 | Δ *trans* + |
| + | $R_P > R_H$ | + | Δ *cis* + *trans* + |
| + | $R_P > R_H$ | - | Δ *cis* × *trans* + |
| - | - | 0 | Δ *trans* - |
| - | $R_P < R_H$ | + | Δ *cis* × *trans* - |
| - | $R_P < R_H$ | - | Δ *cis* + *trans* - |

**Supp. Table 1**: Geometric assignments of gene regulatory differences. Assignments of gene expression differences to *cis, trans, cis + trans* or *cis* × *trans* regulation can be made based on the sign of values of $R_P - R_H$ (log2 of parental fold change minus log2 of hybrid fold change) and $R_H$ (log2 of hybrid fold change). Possible value pairs and the corresponding regulatory assignments are enumerated above.

### Supplementary figures

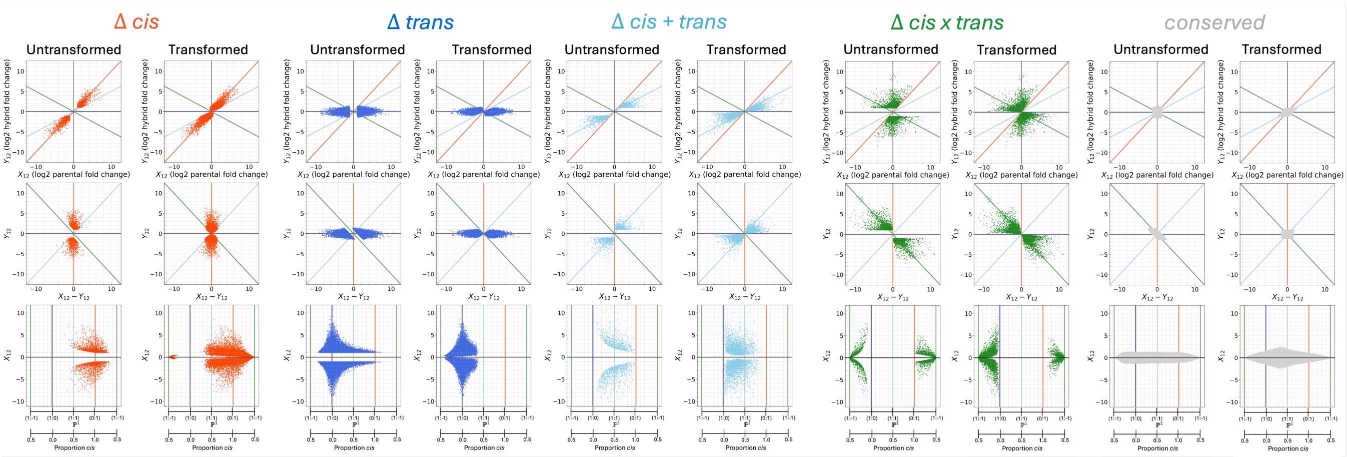


**Supp. Fig. 1**: The points in Figures 2A and 2C separated by regulatory assignment (color). Across all assignments, n = 285,777 gene-cross combinations from 179 unique yeast parent crosses are shown (gene expression data from Tsouris et al., 2024.)

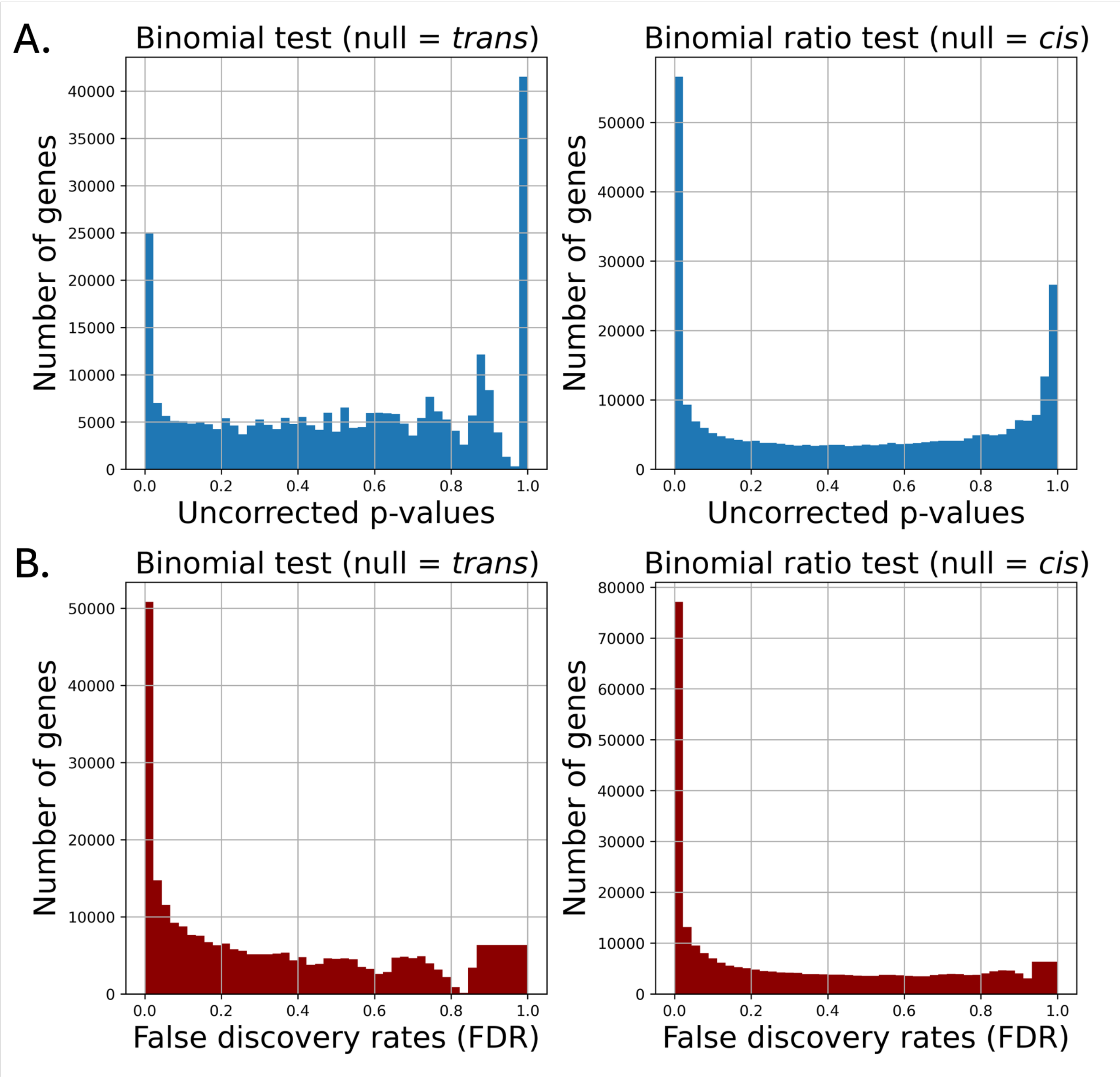


**Supp. Fig. 2**: A) Distribution of p-values for the n = 285,777 yeast gene-cross combinations (Tsouris et al., 2024). B) Distribution of false discovery rates for the same calculated from raw p-values using the Benjamini-Hochberg procedure (Benjamini and Hochberg, 1995).

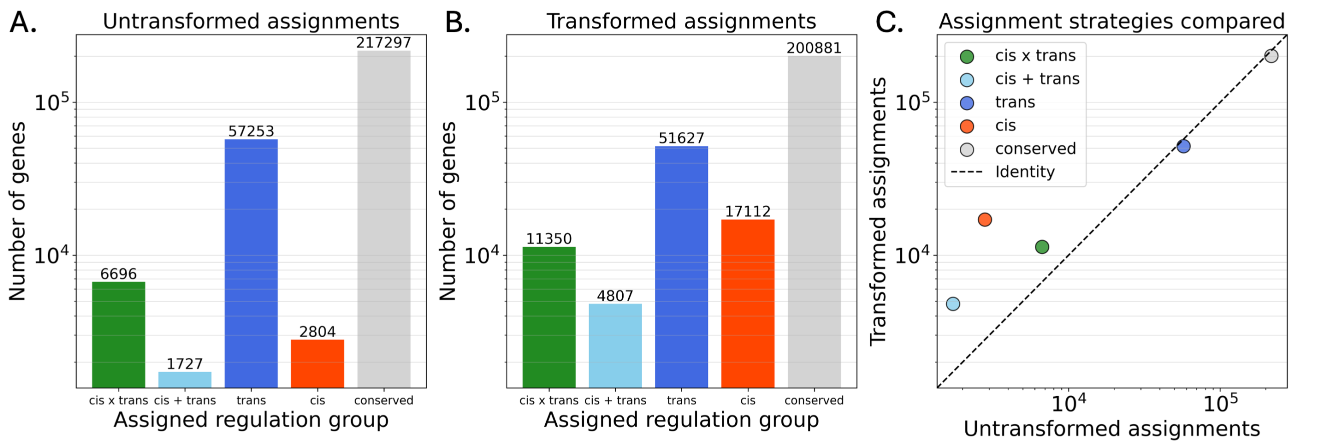


**Supp. Fig. 3**: Comparison of regulatory assignments for n = 285,777 gene-cross combinations in yeast cross experiments (Tsouris et al., 2024). A) The number of genes assigned to designated regulatory categories from Tsouris et al., 2024, and B) our method's assignments. C) Comparison of the number of genes per regulatory category between the previous method (untransformed assignments, x-axis) and our hypothesis testing framework (transformed assignments, y-axis).

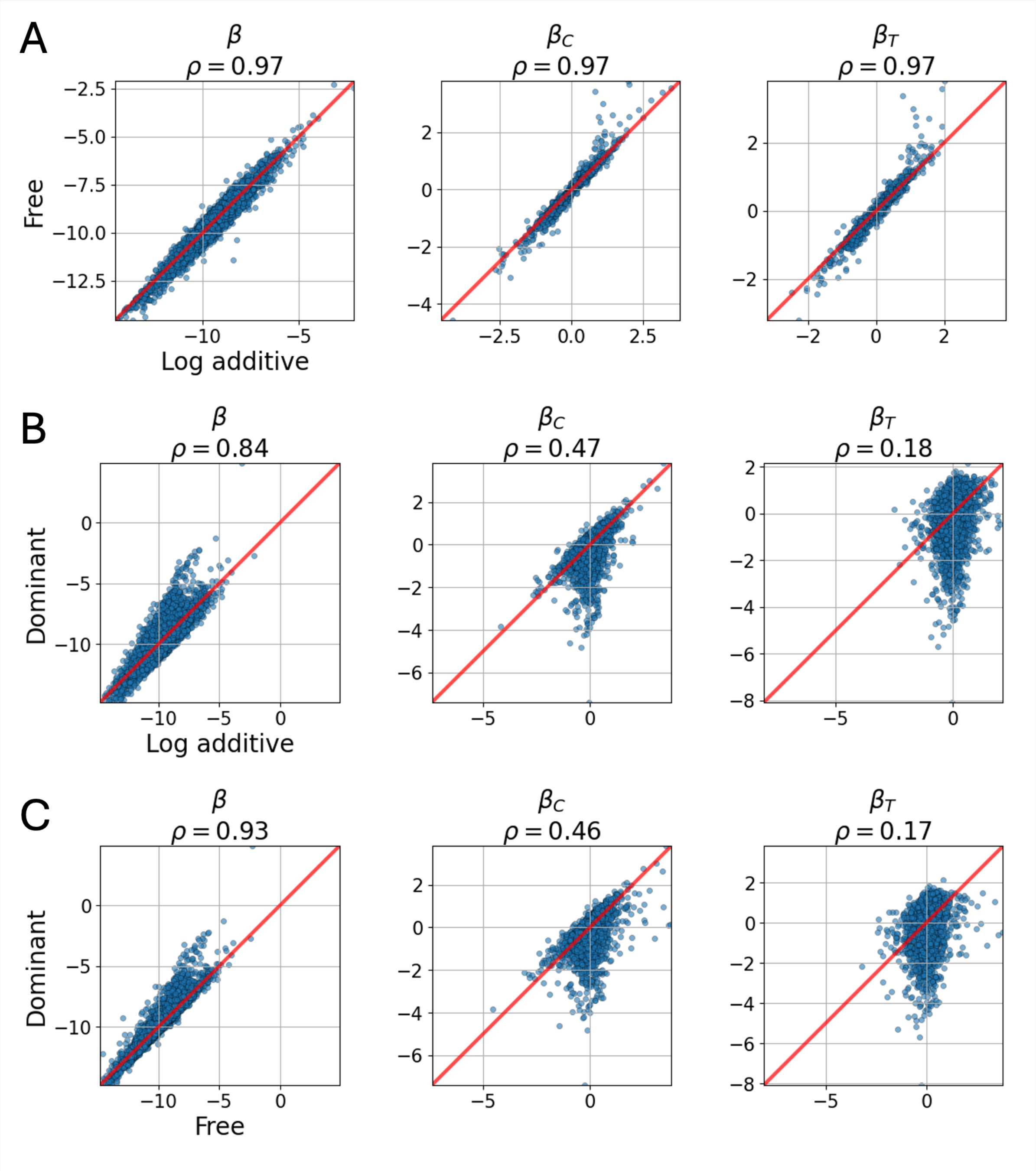


**Supp. Fig. 4**: Comparison of generalized linear model weights fit to BAT tissue samples from mice reared in cold tissues (n = 6 parental strain one, n = 6 parental strain two, n = 6 hybrid crosses) under different model hypotheses (see Methods). $\beta$ is the intercept shared between all samples, $\beta_C$ captures the *cis* effect unique to the non-reference allele, and $\beta_T$ captures the *trans* effect common to hybrids and one parent (see Methods). Pearson correlation coefficients ($\rho$) are displayed for each subplot. A) Log additive model vs. free model, B) log additive vs. dominant model, C) dominant vs. free model. Data from Ballinger et al., 2023.

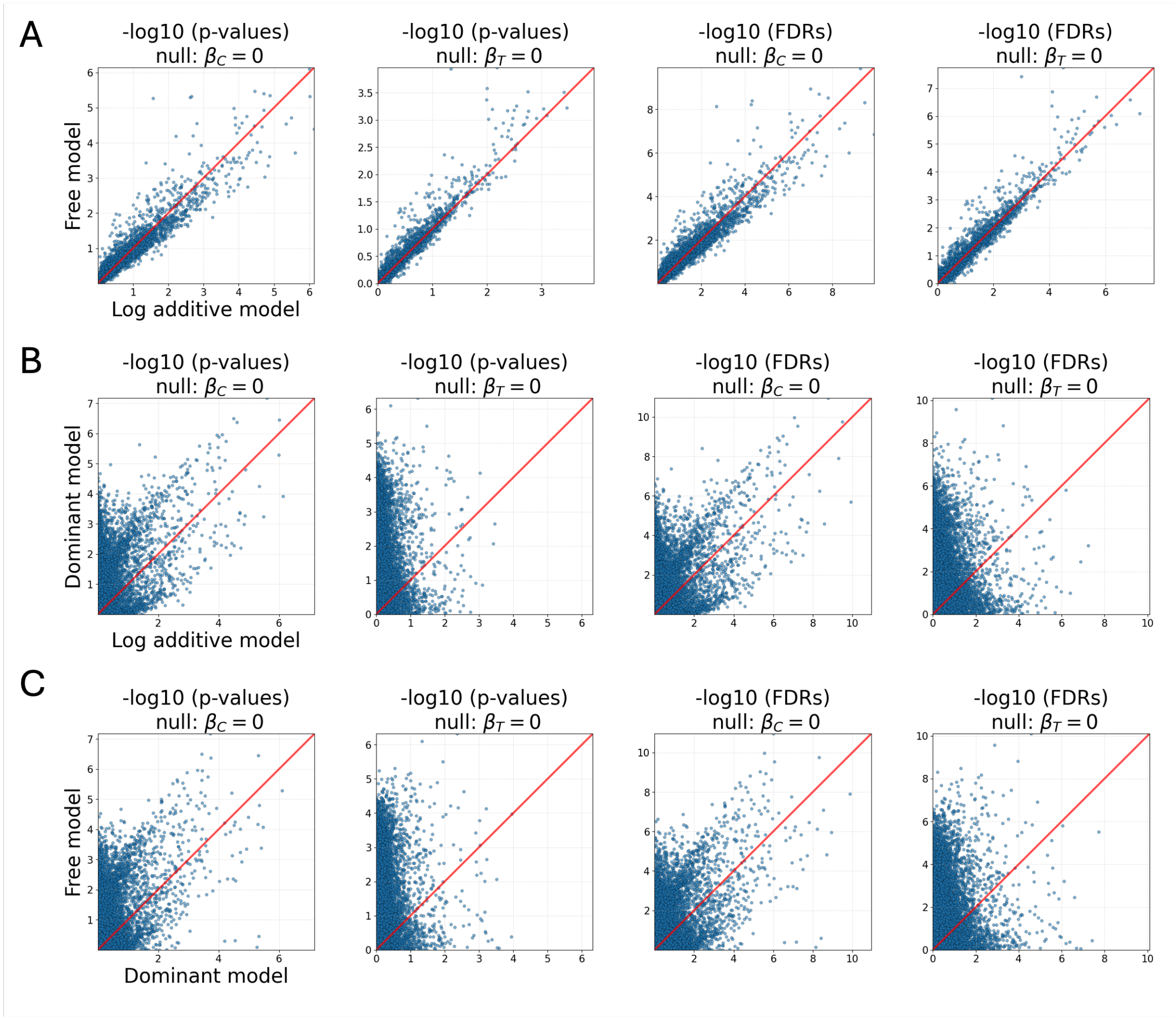


**Supp. Fig. 5**: Comparison of generalized linear model significance values, both raw p-values and Benjamini Hochberg corrected false discovery rates (Benjamini and Hochberg, 1995) for the null hypotheses of no *cis* ($\beta_C = 0$) and no *trans* ($\beta_T = 0$) regulation (see Methods). Comparisons of A) log additive and free models, B) log additive and dominant models, and C) log additive and free models. The models were fit to BAT tissue samples from male mice in reared in the cold (n = 6 parental strain one, n = 6 two, and n = 6 for hybrid crosses with n = 5,970 genes). Data from Ballinger et al., 2023.

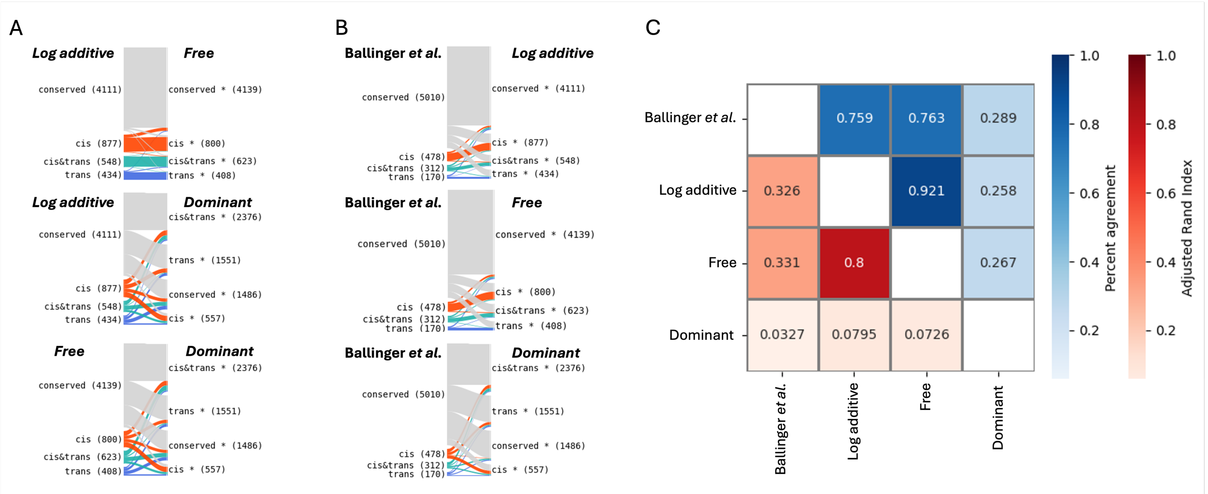


**Supp. Fig. 6**: Gene regulatory assignments made using three different models (see Methods) fit to BAT tissue samples from male mice in reared in the cold (n = 6 parental strain one, n = 6 two, and n = 6 for hybrid crosses for n = 5,970 genes). Data from Ballinger et al., 2023. A) Alluvial plots showing genes categorized as a particular regulatory group in one generalized linear model versus another. B) Alluvial plots showing genes categorized as a particular regulatory group using the previous published method (Ballinger et al., 2023) versus the generalized linear model approach. C) Percent agreement (number of genes classified the same) between models and previous method (blue) and Adjusted Rand Index between model classification (red).

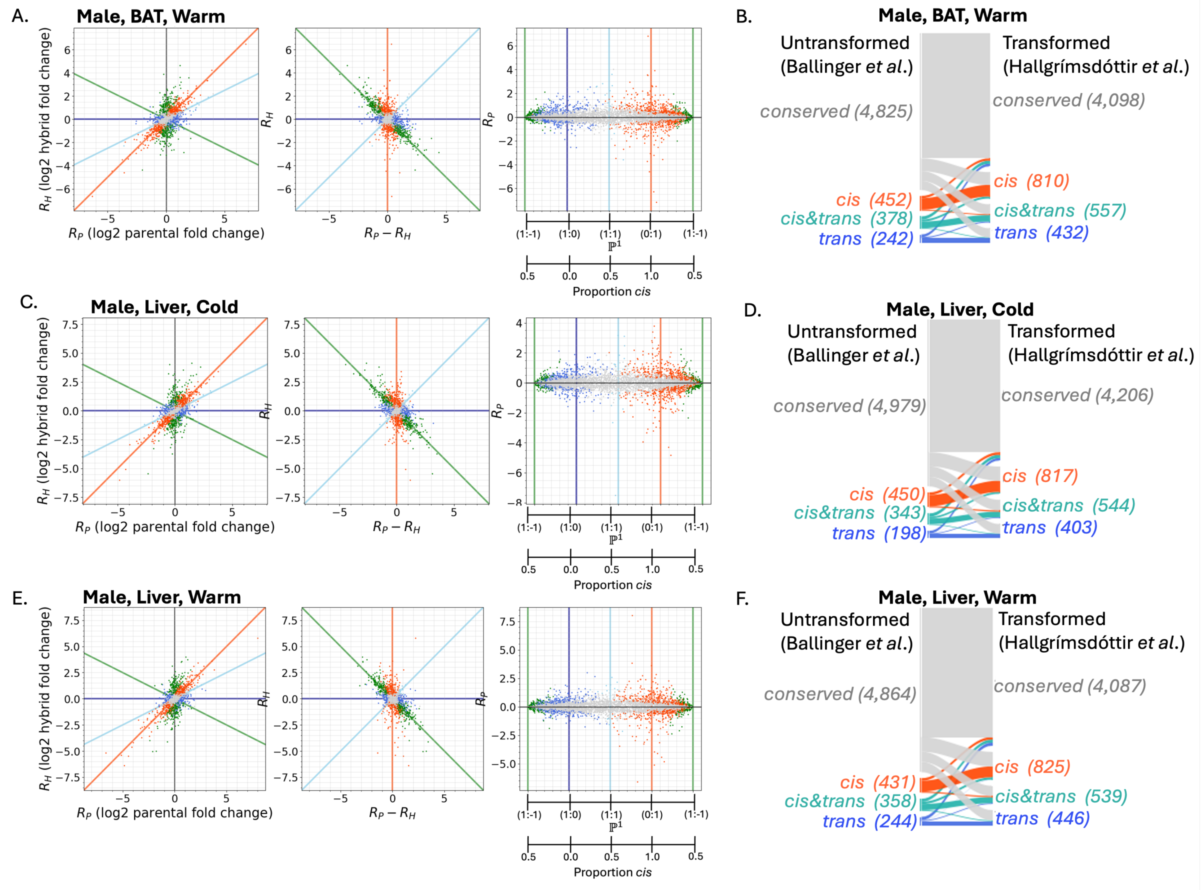


**Supp. Fig. 7**: A) Predictions using a generalized linear model (log additive hypothesis, see Methods) fit to brown adipose tissue (BAT) samples taken from 6 male mice each from two wild-derived inbred lines of house mice (one strain from New York, USA, the other from Brazil) and 6 male mice from a cross between the two lines (Ballinger et al. 2023). All mice were raised in a warm environment (21℃). Log2 of ratio of parent one to parent two counts ($R_P$) plotted against the log2 ratio of parent one allele to parent to allele in hybrids ($R_H$), transformed ratios, and calculated proportion *cis.* The points (N = 5,897) are colored by transformed assigned regulatory category as described in Methods. B) The number in each regulatory assignment from the original study (untransformed), and new assignments (transformed). C) and D) as in A) and B), respectively, but using liver samples taken from mice raised in cold conditions (5℃, 5,970 genes). E) and F) as in A), and B) but using liver samples from mice raised in warm conditions (21℃, 5,897 genes).

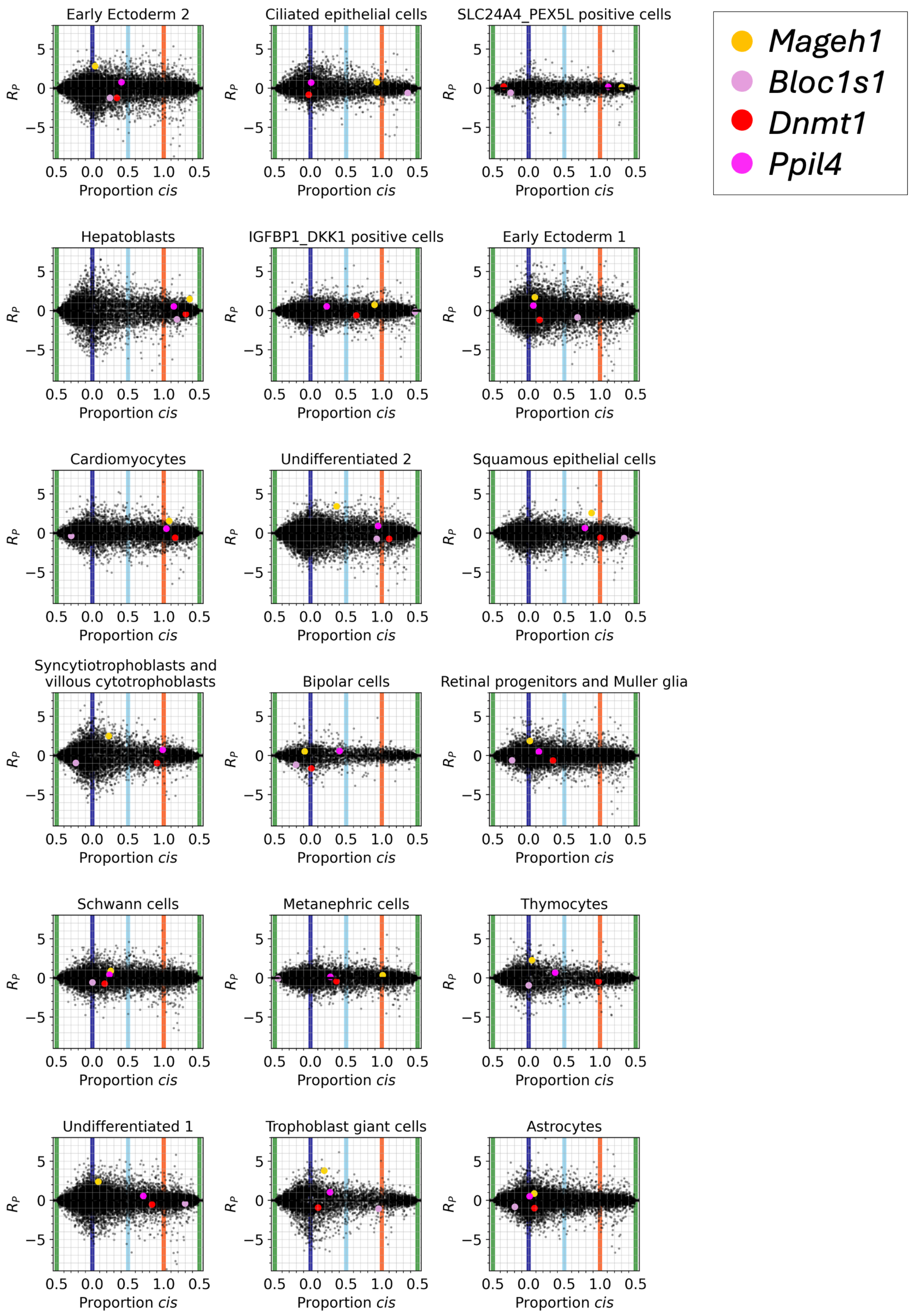


**Supp. Fig. 8**: Proportion *cis* per gene (n = 14,487) separated by cell type calculated using log2 parent and hybrid ratios from gene expression data from human-chimpanzee cell lines (Barr et al., 2023). Results for 18 of the 72 reported cell types are shown. Genes displayed in Main Fig. 3B are colored as indicated in the legend.

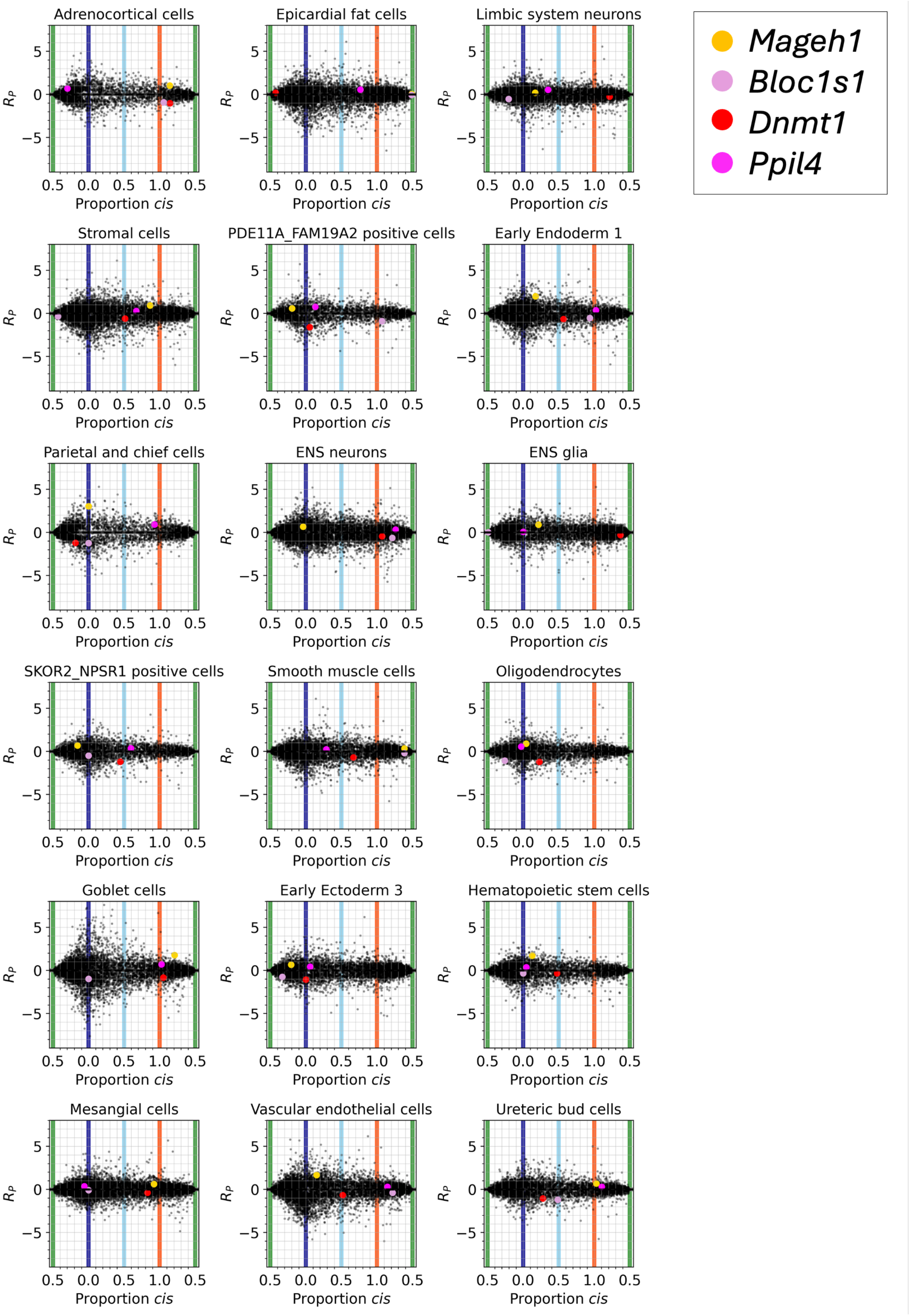


**Supp. Fig. 9**: Proportion *cis* per gene (n = 14,487) separated by cell type calculated using log2 parent and hybrid ratios from gene expression data from human-chimpanzee cell lines (Barr et al., 2023). Results for 18 of the 72 reported cell types are shown. Genes displayed in Main Fig. 3B are colored as indicated in the legend.

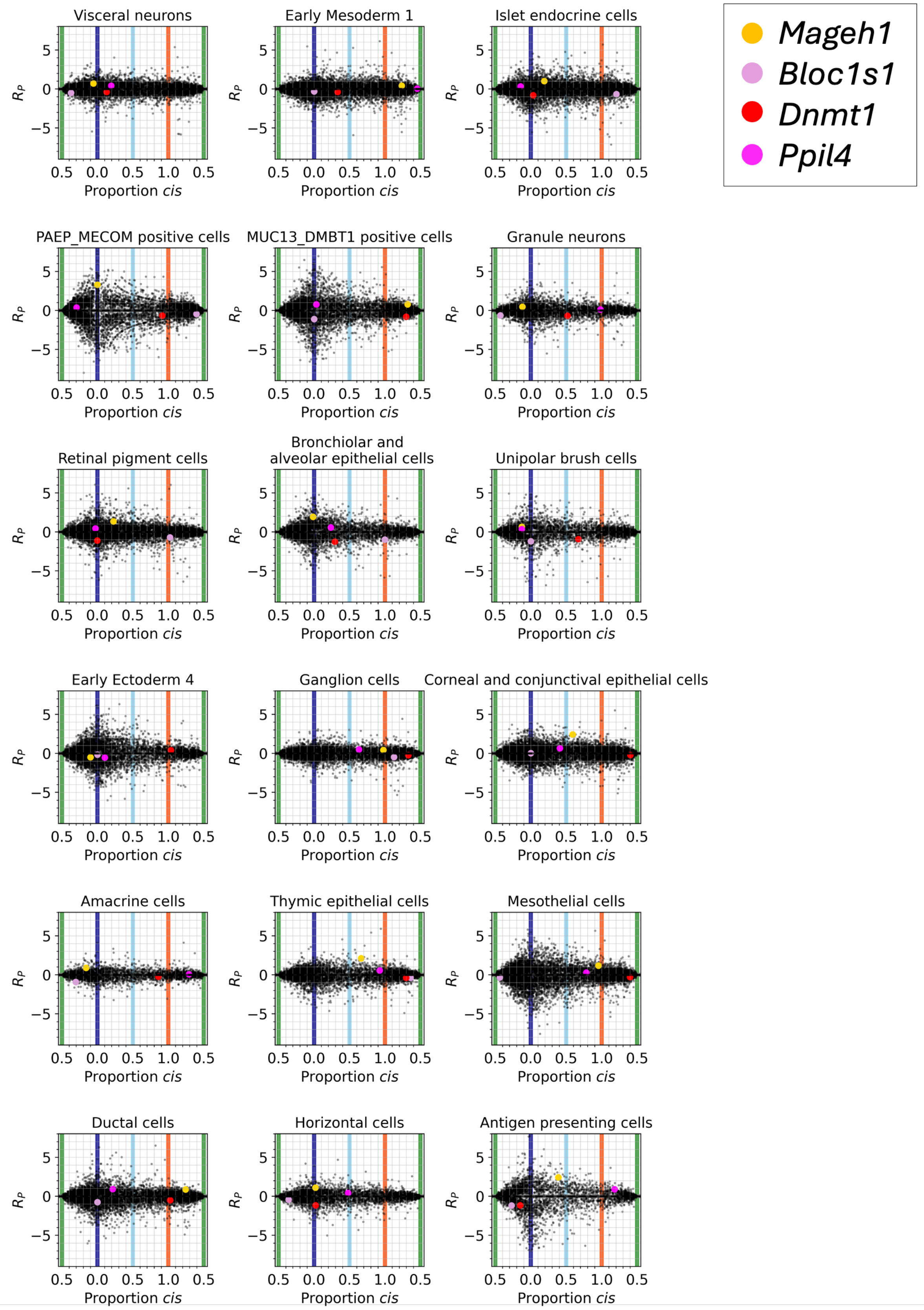


**Supp. Fig. 10**: Proportion *cis* per gene (n = 14,487) separated by cell type calculated using log2 parent and hybrid ratios from gene expression data from human-chimpanzee cell lines (Barr et al., 2023). Results for 18 of the 72 reported cell types are shown. Genes displayed in Main Fig. 3B are colored as indicated in the legend.

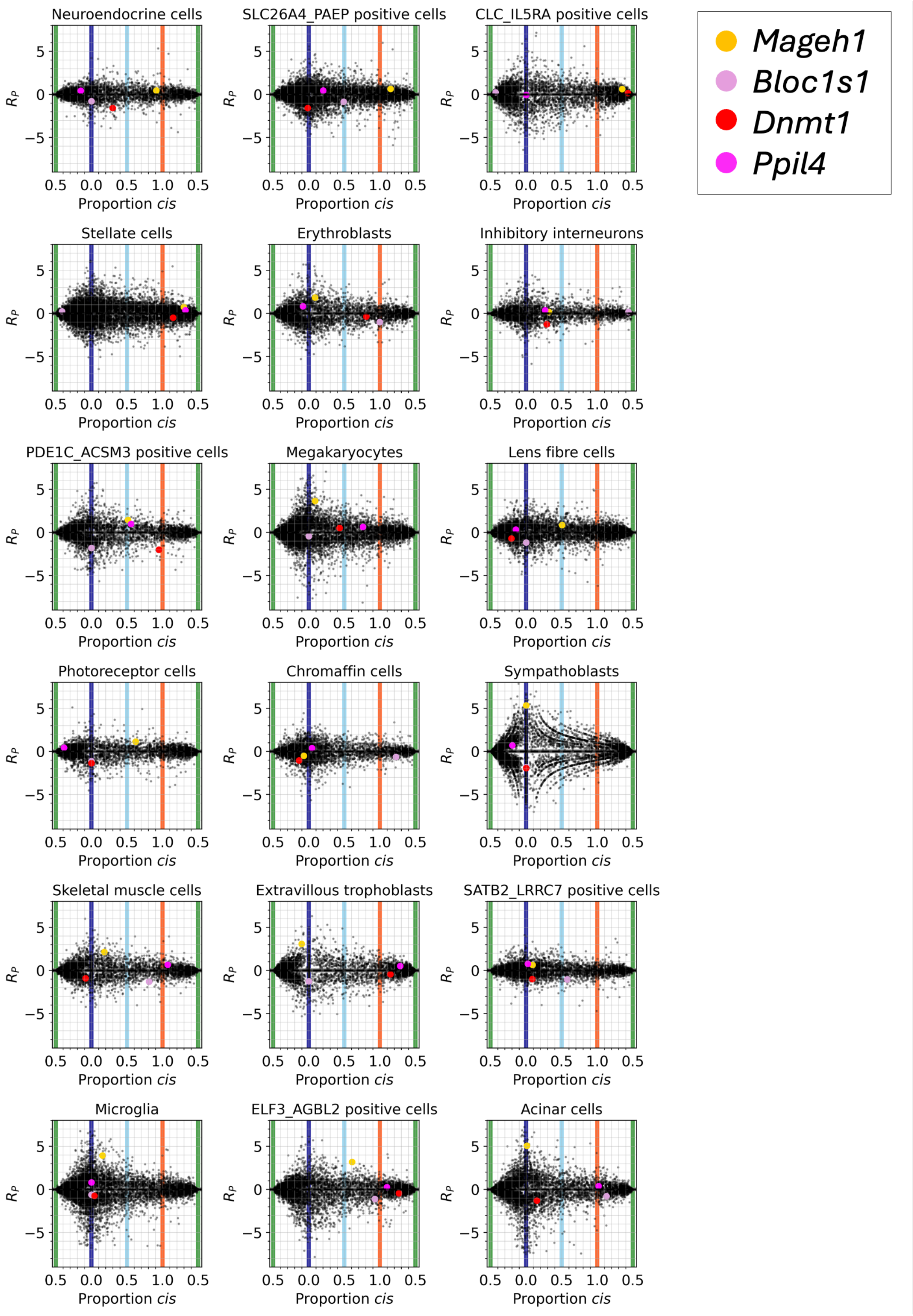


**Supp. Fig. 11**: Proportion *cis* per gene (n = 14,487) separated by cell type calculated using log2 parent and hybrid ratios from gene expression data from human-chimpanzee cell lines (Barr et al., 2023). Results for 18 of the 72 reported cell types are shown. Genes displayed in Main Fig. 3B are colored as indicated in the legend.

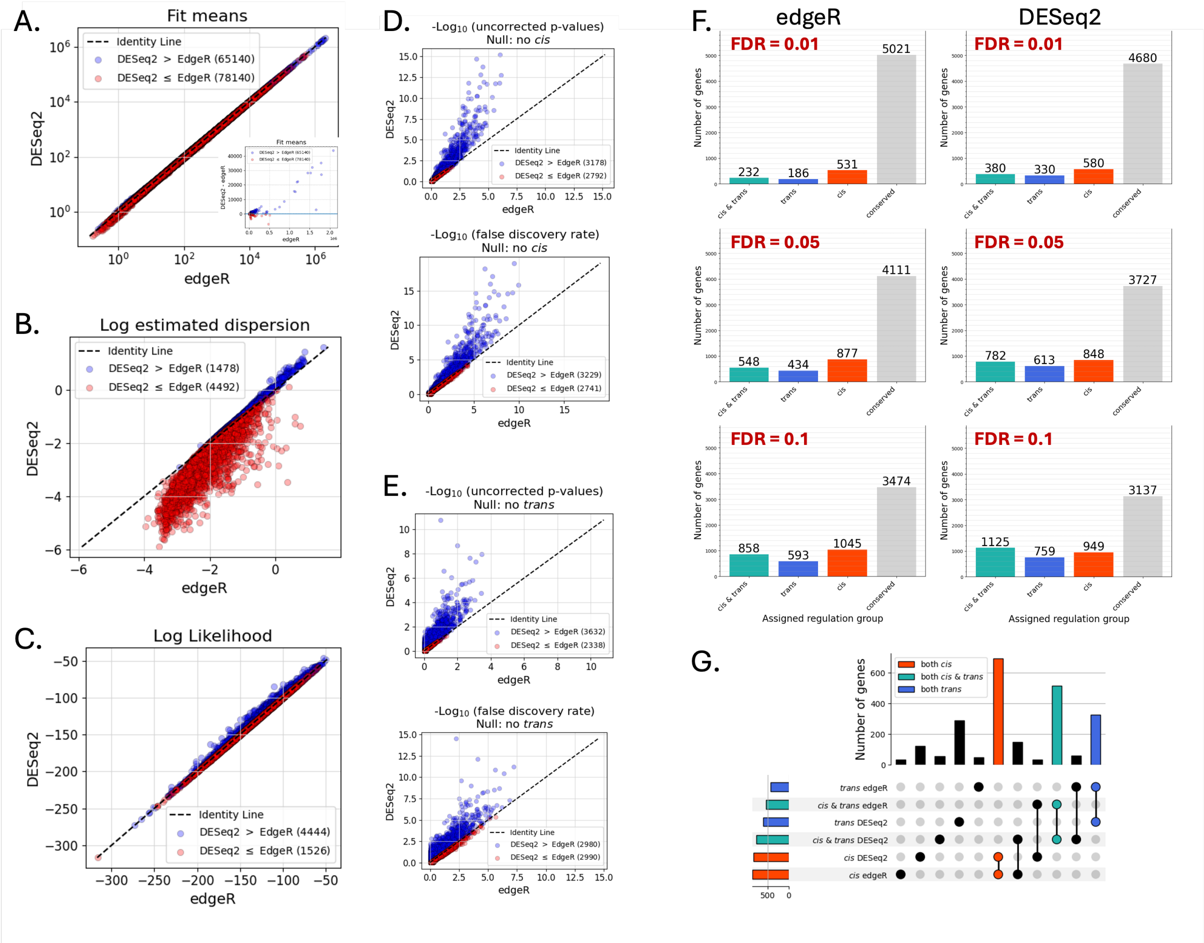


**Supp. Fig. 12**: A comparison of edgeR (Robinson et al., 2010) and DESeq2 (Love et al., 2014) on brown adipose tissue (BAT) samples taken from mice raised in a cold environment (Ballinger et al., 2023). In A)-E), values which are larger in DESeq2 are plotted in blue and values larger in edgeR are plotted in red. A) Model fit means for 5,970 genes are nearly identically between edgeR and DESeq2, but B) DESeq2 dispersion estimates per gene are lower (log of dispersions plotted), which results in C) higher log likelihood values for data under DESeq2 fits than edgeR. D) For the null hypothesis of no *cis* regulation (see Methods), log10 of uncorrected p-values (upper) derived from a likelihood ratio test and false discoveries rates (lower) corrected using the Benjamini-Hochberg procedure (Benjamini and Hochberg, 1995) for DESeq2 vs. edgeR. E) As in D), but for the null hypothesis of no *trans* regulation. F) Assigned regulatory groups (see Methods) using edgeR (left column) or DESeq2 (right column) and different FDR thresholds (0.01, 0.05, and 0.1) show that edgeR classifies more genes as conserved than DESeq2 at the same FDR thresholds. G) Upset plot comparing the genes assigned to *cis, trans* or *cis & trans* for edgeR or DESeq2.
